# Genetic overlap between educational attainment, schizophrenia and autism

**DOI:** 10.1101/093575

**Authors:** Varun Warrier, Richard AI Bethlehem, Daniel H Geschwind, Simon Baron-Cohen

**Author notes:** Corresponding author: Varun Warrier and Simon Baron-Cohen.

## Abstract

**Importance:** The genetic relationship between cognition, autism, and schizophrenia is complex. It is unclear how genes that contribute to cognition also contribute to risk for autism and schizophrenia.

**Objective:** To investigate the interaction between genes related to cognition (measured via proxy through educational attainment, which we call ‘edu genes’) and genes/biological pathways that are atypical in autism and schizophrenia.

**Design:** Genetic correlation and enrichment analysis were conducted to identify the interaction between edu genes and risk genes and biological pathways for autism or schizophrenia.

**Results:** First, edu genes are enriched in a specific developmental co-expression module that is also enriched for high confidence autism risk genes. Second, modules enriched for genes that are dysregulated in autism and schizophrenia are also enriched for edu genes. Finally, genes that overlap between the two above modules and educational attainment are significantly enriched for genes that flank human accelerated regions, suggesting increased positive selection for the overlapping gene sets.

**Conclusion:** Our results identify distinct co-expression modules where risk genes for the two psychiatric conditions interact with edu genes. This suggests specific pathways that contribute to both cognitive deficits and cognitive talents, in individuals with schizophrenia or autism.

**Key Points:** *Question:* How do genes for educational attainment interact with risk genes for autism and schizophrenia?

*Findings:* We show that genes for educational attainment (edu genes) are significantly likely to be mutated in autism and intellectual disability. We further show that edu genes also interact with co-expression modules that are associated with autism or schizophrenia and are enriched for differentially expressed genes in autism or schizophrenia. Finally, we identify that the enrichment between risk genes for autism and schizophrenia and human accelerated regions are driven, in part, by their overlap with edu genes.

*Meaning:* Edu genes interact with schizophrenia and autism risk genes in specific pathways, contributing to both cognitive deficits and talents.

## Introduction

Autism and schizophrenia are complex psychiatric conditions with considerable phenotypic heterogeneity and significant heritability. Both conditions are genetically highly heterogeneous, with common and rare variants implicated in their aetiology. Approximately 40% of individuals with autism have intellectual disability defined as IQ < 70^1^ and as a group, individuals with autism show deficits in theory of mind^2^ and central coherence^3^. In contrast, some studies show superior functioning in domain specific aspects of cognition in individuals with autism, including superior attention to detail^4^, stronger systemizing^5,6^, hypercalculia^7,8^ and savantism^9^. Furthermore, elevated rates of autism and autistic traits have been reported in those working in science, Genetic correlation was performed using technology, engineering and maths (STEM) fields^10,11^. The relationship between IQ/cognitive ability and schizophrenia is also complex, but in a different way. Cognitive difficulties are also observed in individuals with schizophrenia^12^. Other studies have identified a link between schizophrenia and creativity^13^, with polygenic risk scores for schizophrenia predicting creativity. Clearly, both autism and schizophrenia have a complex genetic link with cognition and IQ. This study aims to understand this set of genetic relationships.

There is an established positive genetic correlation between autism and different measures of cognitive attainment such as years of education^14^ and a derived measure of general cognition in children^15,16^. Polygenic risk score analysis has identified significant risk between general cognitive ability and autism^17^ and creativity and schizophrenia^13^. In addition, conditional False Discovery Rate based analysis using genome-wide association study (GWAS) for educational attainment has identified common variants associated with schizophrenia^18^. On the other hand, individuals with autism who are co-morbid for intellectual disability (ID) have a significantly higher rate of de novo variants than individuals with autism who have average or above average IQ^19^.

It is unclear, at a genetic and transcriptional level, how exactly genes for educational attainment interact with genes that contribute to risk in autism and schizophrenia. Two broad, non-mutually exclusive hypotheses might elucidate the complex relationship between cognition/educational attainment and the two psychiatric conditions at a biological level. First, mutations in genes related to educational attainment (what we call ‘edu genes’) might lead to autism and schizophrenia co-morbid with intellectual disability and other cognitive deficits. In other words, there may be an upstream convergence in shared genetic mutations. Second, edu genes might interact with risk pathways for schizophrenia and autism in specific transcriptional co-expression modules, leading to an interaction in common downstream biological pathways. This could lead to increased risk for the psychiatric conditions with preserved or above average cognitive ability.

A few studies have identified a role for human accelerated regions (HARs) in autism ^20^, schizophrenia^21^, and cognition^20^. HARs are conserved regions of the genome that show human-specific divergence at an accelerated pace, thought to indicate positive selections in humans. Genes in these regions may thus have contributed to human-specific traits. Indeed, mutations in genes that lie in HARs affect cognition and contribute to autism^20^. Furthermore, schizophrenia-associated loci are enriched in genes that lie in or close to HARs^21^. It is unclear why HARs are enriched for genes that contribute to both these conditions. One explanation is that genes that lie in or near HAR contribute pleiotropically to both cognition and risk for schizophrenia or autism. This might explain the positive selection in these regions in spite of their association with autism and schizophrenia, two conditions with reduced fecundity^22^. Thus, we hypothesize that genes flanking or interacting with HARs may contribute to the genetic and phenotypic overlap seen between cognition and the two psychiatric conditions.

In this study, we examine the relationship between edu genes and genes and transcriptional co-expression modules implicated in autism or schizophrenia. We identify a specific gene co-expression module in the developmental transcriptome of the human cortex that is enriched for edu genes and genes that are frequently mutated in autism. We further identify co-expression networks that are enriched for differentially expressed genes in schizophrenia and autism and are also enriched for edu genes. Finally, we also demonstrate that the overlapping genes between edu attainment and the co-expression modules enriched for differentially expressed genes for autism or schizophrenia are also enriched for HARs. A schematic diagram of the study design is provided in Figure 1.

**Figure 1:**
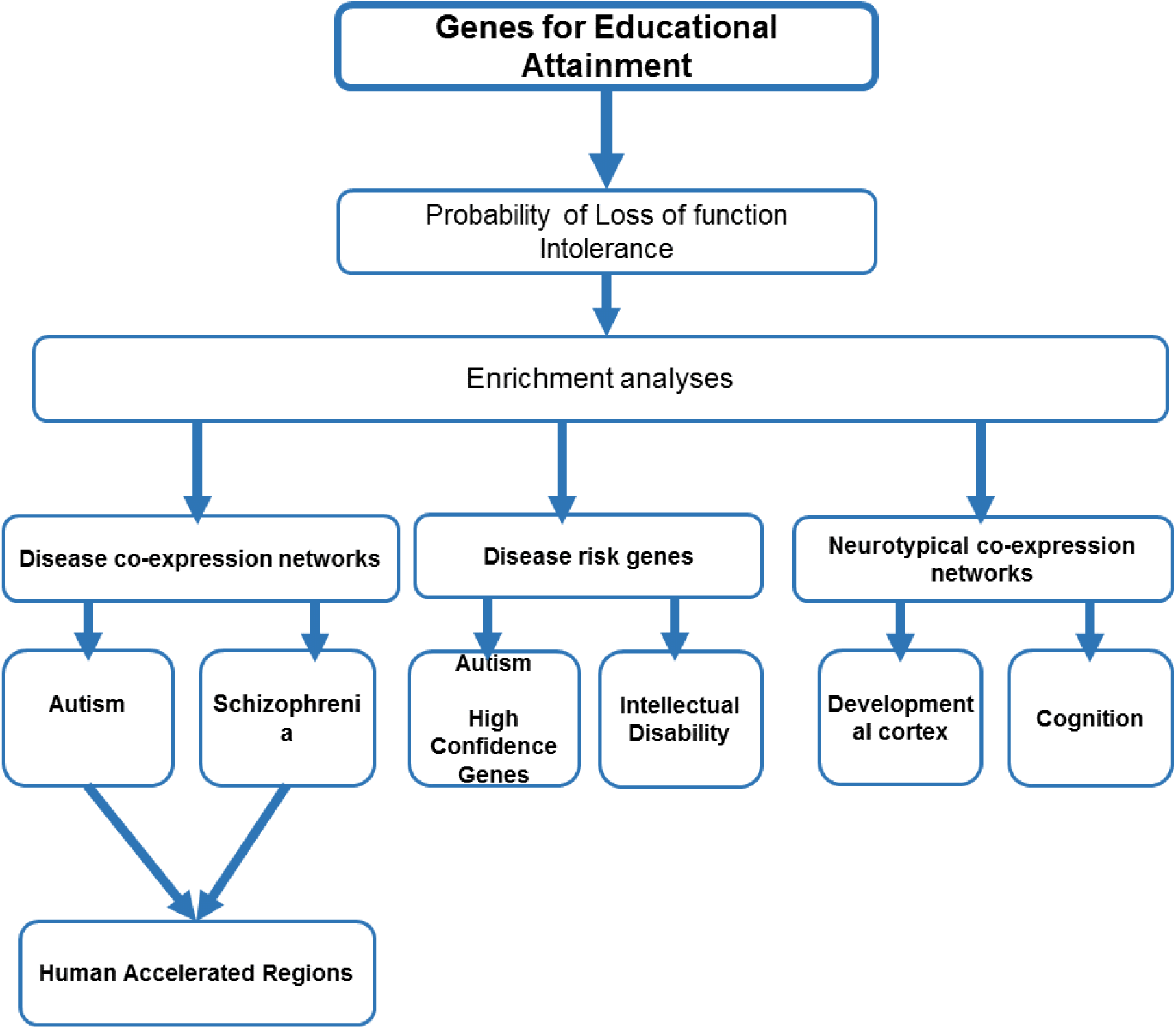
Schematic diagram of the study protocol. Schematic diagram of the study protocol. We first tested for probability of loss of function intolerance for the edu genes and compared with all brain expressed genes. We next tested the enrichment of edu genes in various previously identified weighted gene co-expression modules for autism, schizophrenia, cognition and the developmental cortical transcriptome. We also checked for overlap between edu genes and high-confidence autism genes and intellectual genes. We finally investigated if the enrichment in the autism and schizophrenia co-expression module was guided by genes flanking human accelerated regions.

## Methods

### Genes

We used a list of genes previously identified using DEPICT^23^ for educational attainment^24^ (edu genes). These genes were identified using independent GWAS loci with P < 1×10^−5^. SNPs were defined as being independent if they had an r^2^ < 0.1, and if they were 500kb apart. The genes were identified using a sample of 405,072 participants as previously described by Okbay et al., (2016)^24^. As a control, we also used genes previously identified using DEPICT for height (641 genes)^25^ identified using independent GWAS loci with P < 5x10^−8^ (height genes). DEPICT uses data gathered from 77,870 microarray experiments expressing levels of 19,997 genes covered by the microarray platforms. To investigate enrichment in autism and intellectual disability, a list of genes implicated in autism (henceforth, high confidence autism genes) was obtained from Sanders et al (2015)^26^ and a list of genes implicated in intellectual disability (henceforth, ID genes) from Vissers et al, (2016)^27^.

#### Brain-expressed genes

We identified a list of genes expressed in the brain using expression data from the Gene Tissue Expression portal (GTEx v6)^28^. We downloaded the list of median gene expression per tissue and identified a list of genes expressed in any of the 11 brain tissues (amygdala, frontal cortex, cortex, cerebellar cortex, cerebellum, anterior cingulate cortex, caudate, hippocampus, hypothalamus, nucleus accumbens and putamen). Genes expressed with a median RPKM > 1 in any of the 11 tissues were considered brain-expressed genes. We identified 15,395 brain-expressed genes.

#### Human accelerated region genes

We identified a pre-compiled list of genes found in the human accelerated regions (HAR) using data in multiple different publications^20^. We used the list of genes closest to HARs (HAR genes), and a separate non-independent list of genes that interact with HARs, identified through HiC and Chia-PET experiments (HAR interacting genes)^20^. There is a considerable overlap between the two gene lists. This is not surprising as, according to Doan et al., (2016)^20^, more than 42% of the HARs interact with the flanking promoters. While HAR interacting genes provide evidence for the genes that are influenced by HAR when compared to HAR genes, we consider this list to be preliminary as interaction between HARs have not been investigated in multiple tissues and across multiple time points.

### Co-expression modules

#### Cognition

To test the enrichment of edu genes in networks associated with general cognition, we used two previously identified modules for cognition^29^: M1 (renamed M_c_1) and M3 (renamed M_c_3). Initial weighted gene co-expression networks were constructed using 122 frozen hippocampus samples that were surgically removed from patients with temporal lobe epilepsy. Significant modules within these weighted networks were identified (empirical P ≤ 0.002) after ensuring that co-expression patterns were unrelated to epilepsy, co-expression modules were preserved in rodent hippocampus and preserved across different human neural tissues by comparing the average topological overlaps of the of the genes in each module between different tissues.

#### Autism, schizophrenia, and the developmental transcriptome

We used previously identified gene co-expression modules for both autism^30,31^ and schizophrenia^32^. Due to the relatively small sample sizes of existing transcriptome projects, we used all nominally significant differentially co-expressed modules for both autism and schizophrenia compared to controls. For autism, we used modules mod1 (renamed M_A_1), mod5 (renamed M_A_5), mod6 (renamed M_A_6) from Gupta et al., (2014)^30^. Briefly, RNA sequencing was performed for multiple cortical tissues obtained from the Autism Tissue Programme. In total, there were 47 (32 unique individuals) autism samples, and 57 (40 unique individuals) control samples. Weighted gene co-expression network analyses (WGCNA) were performed for 13,443 genes combining both cases and control samples. Modules differentially co-expressed for autism were identified using a linear mixed regression framework for case-control status. As a validation, we also used three gene co-expression modules that are associated with autism and are enriched for significantly downregulated genes in the autism post-mortem cortex, identified using a larger dataset^31^. These modules are: M4 (renamed CTX-M_A_4), M10 (renamed CTX-M_A_10), and M16 (renamed CTX-M_A_16). All three modules significantly overlap with the M_A_1 module^31^. Briefly, rRNA-depleted RNA sequencing and WGCNA was performed using 82 cases and 81 control cortex samples from 47 individuals with autism and 44 controls. Co-expression networks were constructed from a total of 16398 genes and lncRNAs^31^. We also used the list of differentially expressed genes in the autism cortex identified after FDR correction from Parikshak et al., (2016)^31^.

For schizophrenia we used data from Fromer et al., (2016)^32^. RNA was extracted and sequenced from the dorsolateral prefrontal cortex from 258 schizophrenia cases and 279 control subjects from the University of Pittsburgh, University of Pennsylvania, and Mount Sinai Hospital brain banks. Co-expression networks were constructed for the control samples separately with a total of 16,423 genes using WGCNA. Modules implicated in schizophrenia were defined as those that harboured an excess of differentially expressed genes for schizophrenia. Four control modules were significantly enriched for differentially expressed schizophrenia genes: M2C (renamed M_S_2), M9C (renamed M_S_9), M11C (renamed M_S_11), and M13C (renamed M_S_13). Three additional control modules were nominally significant for enrichment of differentially expressed schizophrenia genes: M14C (renamed M_S_14), M27C (renamed M_S_27) and M30C (renamed M_S_13). We also used a list of differentially expressed genes in the schizophrenia cortex identified after FDR correction from Fromer et al., (2016)^32^.

We tested all three modules in the autism discovery and validation datasets and all seven modules in the schizophrenia dataset for enrichment of genes associated with educational attainment. In addition, we also utilized co-expressed modules identified using gene-expression datasets of the human developmental transcriptome from Parikshak et al., (2013)^33^. These modules were identified using spatio-temporal expression data from the BrainSpan dataset^34^ and provide insight into the temporal window of this potential modular gene enrichment. We did not include Module 7 in the analyses, as the genes in the module were not highly co-expressed with each other^33^.

### Statistical analyses

Genetic correlation was performed using LDSC^14,35^. We used summary GWAS data for educational attainment^24^ (n = 328,917) downloaded from the Social Sciences Genetic Association Consortium website (http://www.thessgac.org/#!data/kuzq8). GWAS data for schizophrenia (n = 79,845)^36^ and autism (n=10,263) were downloaded from Psychiatric Genomics Consortium website (https://www.med.unc.edu/pgc/results-and-downloads). As a control dataset GWAS data for height (n = 253,288)^25^ was downloaded from the GIANT website (http://www.broadinstitute.org/collaboration/giant/index.php/GIANT_consortium_data_files. Probability of Loss of Function Intolerance score (pLI) was identified for each gene from the Exome Aggregation Consortium web browser (http://exac.broadinstitute.org/)^37^.

Enrichment analyses were performed using hypergeometric tests. For each enrichment pair, the background gene list was identified as the set of total common genes between the two pairs. For example, DEPICT uses expression data from 19,997 genes from 77,840 microarray experiments as the initial input to identify enriched genes and gene-sets^23^. A total of 13,262 genes were used to construct a weighted co-expression network for autism genes^30^, of which 11,963 genes were common between the DEPICT gene list and the autism co-expression gene list. A total of 16,423 genes were used for network construction for schizophrenia^32^, and, in total, there were 14,112 genes common between the DEPICT gene list and the schizophrenia co-expression gene list. For the enrichment with high confidence autism genes and ID, we used a list of all genes in DEPICT. For enrichment with modules identified with cognition, we used a list of genes common to DEPICT and all genes used in network construction using WGCNA, which was a total of 10,727 genes. Further, for enrichment using only brain expressed genes, a total of 12,625 genes were common between the DEPICT gene list and genes expressed in the brain using the GTEx dataset.

All enrichment analyses were performed using Ensembl gene IDs. A full list of genes used for enrichment analysis is provided in the Supplementary File. We report significant enrichments of all hypergeometric tests with FDR adjusted P-values below 0.05. All hypergeometric tests report P-values for probabilities of drawing x or more (or less in case of an underlap) successes, where x is the number of overlapping genes between the two gene sets tested. Hypergeometric test P-values and fold difference (or fold enrichment/fold change, defined as the ratio of observed to expected overlap) were calculated using: http://nemates.org/MA/progs/overlap_stats.html. To test the specificity of all significant enrichments between educational attainment and the trait of interest, we used height as a negative control.

To identify multi-set enrichment we used the R package ‘SuperExactTest’^38^. We computed 1-tailed P-values with the hypothesis that we expect to find a significant overlap between the different sets. For each of the multi-set enrichment analyses, we identified background genes as those that were investigated across all different sets. For example, for the 3-way enrichment between the M_A_1 autism co-expression module, edu genes and HAR genes, the background list of genes was the intersection of all genes identified in the transcriptional analyses of the autism post-mortem tissues, and all the genes covered in DEPICT. Further, we restricted our HAR genes to only those that were covered in the DEPICT and identified in the autism transcriptome.

## Results

### Genetic correlation analyses

Previous studies have identified genetic correlations between educational attainment and autism, but not with schizophrenia. Here, we used the recent, larger educational attainment dataset to conduct genetic correlations between both educational attainment and both schizophrenia and autism. There was a modest but significant positive genetic correlation between autism and educational attainment (r_g_ = 0.28 ± 0.04; P=2.4x10^−11^) (Figure 2). This correlation is considerably lower than the correlation between childhood IQ and autism (r_g_ =0.5 ± 0.11), but similar to the previously calculated correlation between autism and college years (r_g_ = 0.3 ± 0.08; P = 1.5x10^−4^)^14,15^. In addition, there was a small but significant positive correlation between schizophrenia and educational attainment (rg = 0.096± 0.02; P = 5.39x10^−6^) (Figure 2a). Furthermore, we did not identify significant genetic correlations between height and schizophrenia (0.0011±0.01; P = 0.952) or height and autism (r_g_ = −0.02± 0.004; P = 0.47), suggesting height to be an appropriate negative control.

**Figure 2:**
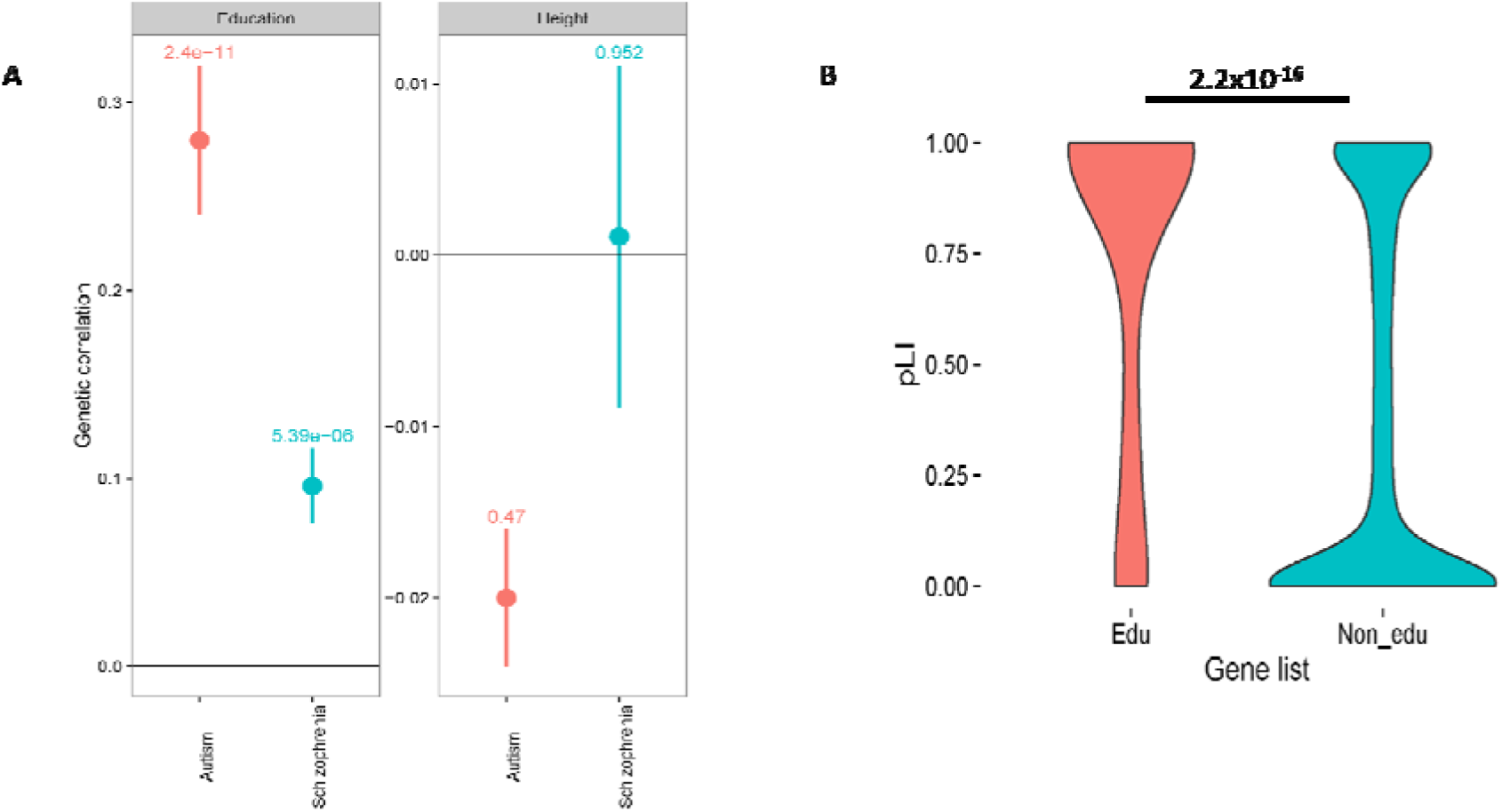
Genetic correlation analyses and distribution of pLI scores for edu genes.

### Genes for education (edu genes) are highly resistant to loss of function

Using data from the Exome Aggregation Consortium (ExAC), we calculated the probability of loss of function intolerance (pLI) for all 146 genes identified for educational attainment. pLI scores were available for 141 of the 146 genes. A pLI => 0.9 is considered to be highly intolerant to loss of function, and is likely to be critical to foetal survival and development. Of the remaining 141 genes, 89 had a pLI => 0.9 (Supplementary Data) that is significant (P = 0.0012; one-sided binomial sign test). In contrast, there were only 19 LoF tolerant genes, suggesting a significant depletion of LoF tolerant genes to educational attainment (P < 0.0001, one-sided binomial sign test). However, brain genes are likely to be more intolerant to other genes. We thus compared the distribution of pLI scores between genes for edu attainment with pLI scores for all brain genes. Edu genes in the brain had higher mean and median pLI scores compared to all brain expressed genes (edu genes vs brain genes; mean: 0.36 and 0.74 respectively, median: 0.08 and 0.98 respectively). However, analysis of the frequency histograms of the pLI scores for both gene sets revealed a nonnormal distribution with heavy tails. Wilcox-rank test identified significantly higher pLI scores for the brain edu genes compared to non-edu brain genes (P < 2.2 x 10^−16^) (Figure 2b). These results suggest that the edu genes, more specifically than simply being brain expressed genes, are unlikely to harbour severe mutations and are likely to be embryonically lethal.

### Edu genes are enriched in a module enriched for general cognitive function

Educational attainment and general cognitive function share a significant, albeit modest, correlation (Proxy correlation: Years of education and Childhood IQ: rg = 0.731± 0.08; P = 2.18×10^−17^)^15^. Genetic correlation between educational attainment and specific aspects of cognitive ability ranged from 0.73±0.04 (for numerical reasoning) to 0.22±0.05 (for reaction times)^39^. A recent study identified two neural modules (M_c_1 and M_c_3) that were enriched for genes implicated in various measures of general cognition^29^. We tested if the edu genes are enriched in the two cognition modules. While we did not identify a significant enrichment on M_c_1, we did identify a significant enrichment in M_c_3 (Fold difference = 3.2; P _FDRadjusted_ = 0.024). We did not identify a significant enrichment for height genes in either of the modules (Supplementary Data). M_c_1 is highly expressed across the cortex, and has a high expression in fetal brain starting at early mid-fetal development (16 ≤ PCW ≤ 19), and increases at birth with the expression persisting postnatally.

### Edu genes are enriched in modules implicated in schizophrenia and autism

We subsequently investigated if these genes have a more downstream effect on schizophrenia and autism, affecting common biological pathways and systems. To investigate this, we used previously identified co-expression modules from adult brain tissues that are enriched for genes that are differentially expressed in autism (three modules: M_A_1, M_A_5, and M_A_6) and schizophrenia (seven modules; M_S_2, M_S_9, M_S_11, M_S_13, M_S_14, M_S_27, M_S_30) (see methods). We identified a significant enrichment for educational attainment genes in M_S_2 (fold difference = 2.9; P_FDRadjusted_ = 8.6 × 10^−8^). In total, there were 34 edu genes that overlapped with the M_S_2 module. In contrast, for the same module, there was no enrichment for genes associated with height (fold difference = 1.1; P_FDRadjusted_ = 0.31). Notably, the M_S_2 module comprises 119 genes that are differentially expressed in schizophrenic post-mortem brain tissue compared to controls. Of this, 117 genes are upregulated in schizophrenia compared to controls. The M_S_2 module is most significant enriched for the gene ontology (GO) term ‘synaptic transmission’. We also identified a modest enrichment for edu genes in the autism M_A_1 module (25 overlapping genes, fold difference = 1.5; P_FDRadjusted_ = 0.049). The M_A_1 module is also enriched for synaptic genes, but is downregulated in autism compared to controls. Again, this enrichment was not significant for height. The M_A_1 module is enriched for genes that contain the GO term ‘synaptic transmission’. To confirm the analysis, we investigated if there is enrichment with Edu genes in three autism co-expression cortical modules identified in a different dataset but that overlaps with the M_A_1 modules (CTX-MA4, CTX-M_A_10, and CTX-M_A_16). All three modules are enriched for genes that are significantly downregulated in autism. We identified a similar modest enrichment of the edu genes in the CTX-M_A_4 (fold difference = 2.61; P_FDRadjusted_ = 0.033) and CTX-M_A_10 modules (fold difference = 2.49; P_FDRadjusted_ = 0.033) and the CTX-M_A_16 (fold difference = 2.57; P_FDRadjusted_ = 0.033) modules. The CTX-M_A_4 and the CTX-M_A_16 module are also enriched for the GO term ‘synaptic transmission’. Taken together, these results suggest that edu genes are enriched in downregulated autism modules that are primarily involved in synaptic transmission.

### Edu genes are enriched for genes implicated in autism and intellectual disability

We next investigated if high confidence autism genes are enriched in educational attainment. We used a list of 65 genes that have been identified using exome and whole genome-sequencing at FDR < 0.1^26^, but restricted it to genes that are both expressed in the brain and are captured in the DEPICT gene list. Enrichment analyses identified 5 common genes between the two datasets, which was higher than expected (fold difference = 8.3; P_FDRadjusted_ = 3.2×10^−4^; Supplementary Data). As a negative control, we tested for enrichment in genes identified for height using DEPICT. We did not identify a significant enrichment (fold difference = 1.7, P = 0.22; Figure 3d). We also investigated if edu genes also contribute intellectual disability, by testing for enrichment using ID. Again, we restricted our analyses only to brain-expressed genes and genes covered in the DEPICT gene list. We identified a significant overlap between intellectual disability genes and edu genes (fold difference = 2; P = 0.013). Interestingly, there was a significant overlap between height and ID genes, though the enrichment was smaller (fold difference = 1.3; P = 0.047). Several genes that contribute to intellectual disability also cause significant changes in height. For example, syndromes like Fragile X and Prader Willi influence both height and intellectual ability^40^.

**Figure 3:**
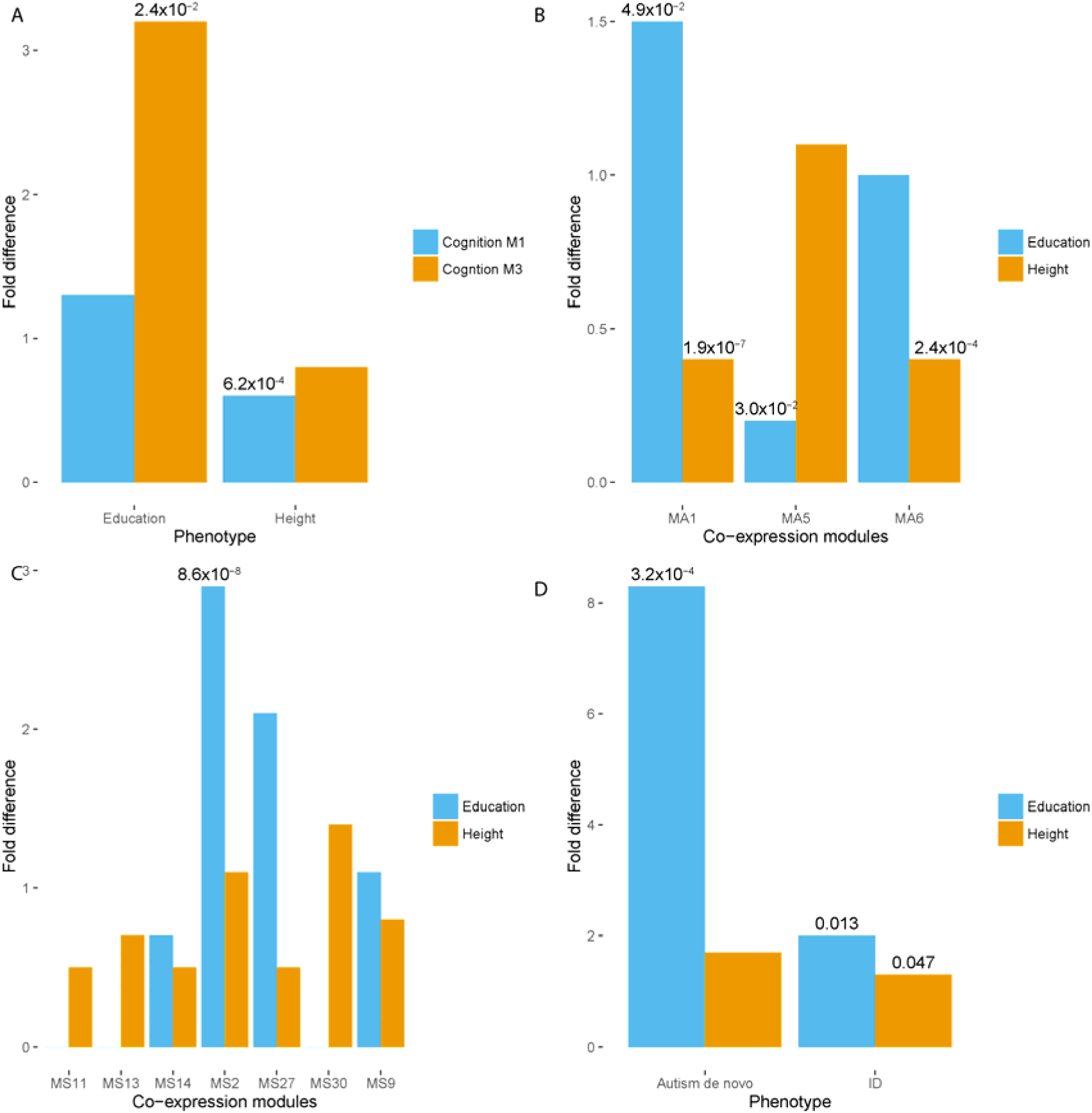
Enrichment analyses. For all the bar charts, y-axes indicate fold difference. A fold difference > 1 is indicative of enrichment, and a fold difference < 1 is indicative of depletion. Significant P-values after FDR correction are provided on top of the bar charts. 3A: Enrichment of edu genes and height genes in the two cognition modules (Mc1 and M_C_3). 3B: Enrichment of edu genes and height genes in three autism co-expression modules (M_A_1, M_A_5 and M_A_6). 3C: Enrichment of edu genes and height genes in the schizophrenia co-expression modules (M_S_11, M_S_13, M_S_14, M_S_2, M_S_27, M_S_30, and M_S_9). 3D: Enrichment of edu genes and height genes in genes implicated through sequencing in autism and intellectual disability (ID).

**Figure 4:**
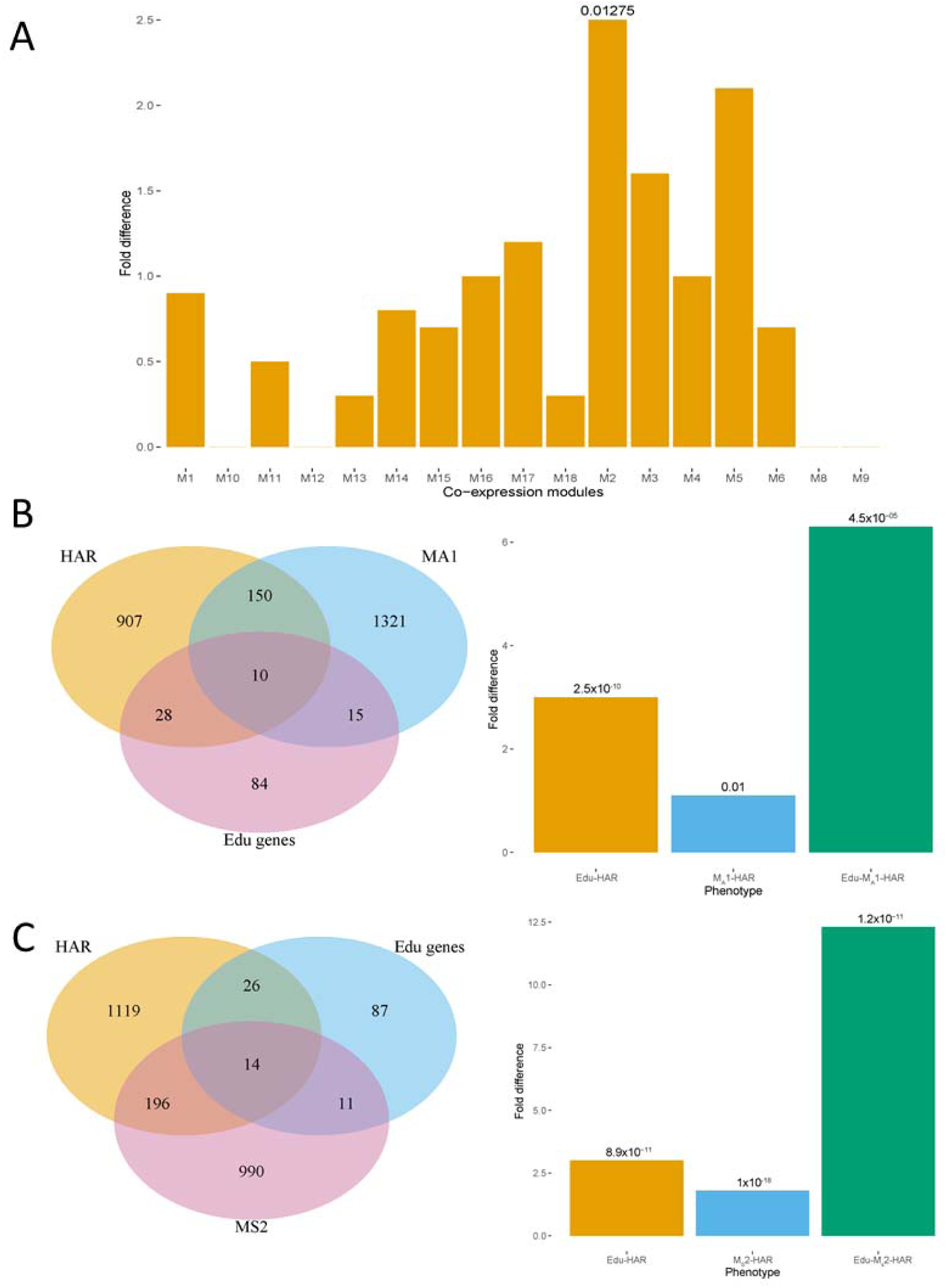
Enrichment in the developmental cortical transcriptome modules and overlap with genes flanking human accelerated regions. 4A: Bar charts showing enrichment of Edu genes in developmental cortical co-expression modules. Y-axis provides fold difference. Significant P-valus after FDR correction provided on top of the graphs. 4B: Venn diagram (left) and bar chart showing overlap and enrichment between Edu genes, autism M_A_1 module and HAR flanking genes. P-values provided on top of bar charts. 4C: Venn diagram (left) and bar chart showing overlap and enrichment between Edu genes, Schizophrenia M_s_2 module and HAR flanking genes. P-values provided on top of bar charts.

### Edu genes are enriched in one module in the developmental transcriptome

We next asked if edu genes are significantly enriched in developmental modules. We utilized modules identified by Parikshak and colleagues (2013)^33^ using the developmental transcriptome data from BrainSpan. We identified a significant enrichment for edu genes in the M2 module (Fold difference = 2.5; P__FDRadjusted_ = 0.01). Interestingly, this module has been reported to be enriched for rare de novo variants in autism, and genes that are implicated in intellectual disability. However, since the Parikshak et al., (2013) paper was published, additional genes have been identified for both autism and ID. We further tested if the M2 module overlapped with high confidence autism genes and ID genes. Indeed this module had significant enrichment for autism genes, (Fold difference = 4.2; P_FDRadjusted = 9.36×10^−5^), but only nominally significant for the expanded ID genes (Fold difference = 1.3; P__FDRadjusted_ = 0.057), supporting the predictive power of these ASD-related gene networks. There was a significant three-way enrichment between genes in the M2 module, high confidence autism genes and edu genes (Fold difference = 122.8; P = 2.2×10^−6^). Three of the 5 overlapping genes between edu genes and the 65 high confidence autism genes lie in the M2 module. This module was not significantly enriched for genes that are differentially expressed in autism brain^31^ (Fold difference = 0.37, P =1) or genes that are differentially expressed in schizophrenia^32^ (Fold difference = 0.6, P__FDRadjusted_ = 0.02).

### Human accelerated region genes are enriched in overlapping gene sets

We reasoned that the genes that overlap between educational attainment and the significant co-expression modules for the two psychiatric conditions might be under stronger selection pressure in humans. To investigate this, we used lists of genes that flank and interact with human accelerated regions (HARs)^20^. HARs are regions of the genes that are conserved in other mammals but are different in humans after they diverged from chimpanzees. These regions are thought to contribute to human specific traits including higher-order cognitive function. We identified a list of genes that interact closely with HARs. To avoid potential confounds, we focused only on brain-expressed genes for all datasets. We conducted a three way-enrichment analysis between brain-expressed Edu genes, brain-expressed schizophrenia M_S_2 genes, and brain expressed HAR genes. We identified a significant overlap between the three sets of genes (Fold difference = 12.31; P = 1.20×10^−11^). This was not significant for height (Fold difference = 1.9; P = 0.05). The overlap between Edu genes and HAR genes was higher (Fold difference = 3.01; P = 8.9x10^−11^) than between M_S_2 and HAR genes (Fold difference = 1.8; P = 1×10^−18^). In addition, the genes overlapping between educational attainment and the MS2 module were significantly more enriched for HAR genes than the non-overlapping M_S_2 genes (OR = 5.4; P = 1.3×10^−7^). However, we did not identify a significant enrichment for HAR genes in the overlapping gene set compared to the non-overlapping edu gene set (OR = 2.1; P = 0.07).

We next investigated the three-way overlap between the M_A_1 module for autism, edu genes and HAR genes. Using only genes expressed in the brain in all three datasets, we identified a significant enrichment between HAR genes, genes in the M_A_1 module, and edu genes (Fold difference = 6.3, P = 4.5×10^−6^). Again, there was no significant enrichment for height (Fold difference = 0.85; P = 0.69). We were able to validate this using the CTX-M_A_4 module (Fold difference = 13.2, P__FDRadjusted_ = 4.89×10^−3^) and the CTX-M_A_10 module (Fold difference = 8.5, P__FDRadjusted_ = 0.036). However, we did not identify a significant enrichment for the CTX-MA16 module (Fold difference = 3.69, P__FDRadjusted_ = 0.23) The enrichment between HAR genes and edu genes was higher (Fold difference = 3.0, P = 2.5×10^−10^) than between M_A_1 and HAR genes (Fold difference = 1.1; P = 0.01). Again, the overlapping genes between edu genes and M_A_1 were significantly more enriched for HAR genes than non-overlapping M_A_1 genes (OR = 6.59; P = 2.4×10^−7^), but the enrichment was non-significant when comparing the overlapping genes with the non-overlapping edu genes (OR = 2, P = 0.12). We also investigated if the overlaps would remain significant if considering only HAR-interacting genes. We did not observe a significant overlap for either of the datasets (Edu-M_S_2-HAR_interacting: Fold difference = 3.9; P = 0.09; Edu-M_A_1-HAR_interacting: Fold difference = 1.2; P = 0.54).

## Discussion

Here, we provide evidence for the interaction between edu genes and risk genes and co-expression networks for autism and schizophrenia. We further show that genes that overlap between the co-expression networks encompassing disease risk and genetic association for educational attainment are significantly enriched for genes flanking human accelerated regions^20^, suggesting that the regulation of these overlapping genes is under positive selection in humans. This overlap with educational attainment likely also explains the previous observations that schizophrenia risk loci are also enriched near HARs^21^. We show that, for autism, both genes that are frequently mutated in autism, and genes that have differential expression in post mortem autism brain, interact with edu genes, but in distinct co-expression modules that represent different biological pathways.

We built on previous research that identified a positive genetic correlation between common variants educational attainment and common variation in two psychiatric conditions – autism and schizophrenia^14,15^. We replicated this correlation using the latest educational attainment GWAS, which has more than twice the number of participants from the previous GWAS on educational attainment. To identify points of convergence between these seemingly disparate phenotypes, we tested two broad hypotheses. The first is that the same genes that are implicated in educational attainment are also more frequently mutated in autism and learning difficulties or intellectual disability (ID) *(the upstream hypothesis).* The second is that educational attainment genes converge in more downstream cellular and molecular pathways *(the downstream hypothesis).* Our results provide preliminary evidence for the involvement of edu genes in both mechanisms.

The significant overlap between edu genes and high confidence autism risk genes is consistent with the observation that approximately 40% of individuals with autism have comorbid ID^1^. This represents the *upstream hypothesis,* where mutations in specific genes are likely to have large effects. Hence, deleterious mutations in edu genes cause more severe phenotypes with marked cognitive deficits such as ID and autism. This is supported by the high PLI scores for edu genes compared to all brain expressed genes. Both edu and high confidence autism risk genes converge on common neural pathway, including synapse development and maintenance. In addition, both sets of genes are active in the fetal brain^24,33^. Our results also show that this enrichment is not specific to autism alone, but can be extended to ID, using genes identified in multiple cohorts.

We further identify a co-expression module (M2) in the developmental transcriptome of the cortex that is enriched for both high confidence, large effect size, autism risk genes^33^ and edu genes, which is supported by a highly significant three-way overlap between edu genes, genes in the M2 co-expression module, and high confidence autism risk genes^33^. Since the M2 module was originally identified with a relatively small number of ASD risk genes^33^, its greater than 4-fold enrichment with subsequently identified risk genes validates the original prediction that such network modules are indeed enriched in ASD risk and can be used to prioritize candidate genes^33^. Notably, this module was not significantly enriched for genes that are differentially expressed in schizophrenia or autism underscoring previous reports that show that differentially expressed genes and high confidence, large effect size, autism risk genes may lie in different co-expression networks^30,33^.

The M2 module is highly preserved in utero and in early infancy and is enriched for genes involved in the GO processes chromatin binding and regulation, and thus is thought to be involved in neural differentiation and axon guidance. Mutations in these early processes likely contribute to global changes in neural architecture leading to more severe phenotypes, and thus, it is not surprising that high confidence autism genes and edu genes converge in this co-expression module. While variants with small effects in these genes contribute to higher educational attainment, more deleterious mutations can have the opposite effect and significantly reduce cognitive function leading to autism co-morbid with ID. To our knowledge, far fewer major risk genes for schizophrenia have been identified with similar statistical support as for ASD^41^, and as such we do not have sufficient power to test for overlap. One previous study has investigated the enrichment of mutated genes in autism, schizophrenia, and ID with educational attainment^24^. However, here we focussed only on the subset of high confidence genes for autism and included an expanded ID gene set.

We also identify an enrichment for edu genes in the schizophrenia M_s_2 and the autism M_A_1, CTX-M_A_4, CTX-M_A_10, and CTX-M_A_16 modules using the largest yet cortical transcriptome dataset for schizophrenia and autism that is publically available. We note that the autism transcriptome datasets are considerably smaller than the schizophrenia dataset, and as such, affords lower power for subsequent analyses. All the modules represent more *downstream processes* involved in the two conditions, and can be thought to reflect both genetic and environmental processes that contribute to the two conditions. The M_S_2 module comprises genes that are upregulated in schizophrenia when compared to controls, whereas the autism modules comprise genes that are downregulated in autism. Notably, the M_S_2, M_A_1, CTX-M_A_4 and CTX-M_A_16 modules are also enriched for genes involved in synaptic transmission, suggesting that they capture similar underlying biological processes. The M_s_2 module was also enriched for common variants associated with schizophrenia^32^. Further, both the overlapping gene sets were significantly enriched for genes flanking HARs, suggesting that they may be under positive selection, maintaining the variants in the general population. Previous research has showed that genetic loci associated with schizophrenia, a condition with reduced fecundity, are enriched near HARs, which are under positive selection. Here we show that this association may be driven by the pleiotropic overlap with educational attainment. We found that, at the level of transcriptional co-expression modules, risk genes for schizophrenia interact closely with genes that contribute to educational attainment. We show that genes that overlap between schizophrenia and edu attainment have higher enrichment for HAR genes than edu genes alone, or the M_s_2 genes alone. We also identify a similar mechanism of enrichment with the autism co-expression modules. This pleiotropy at the level of transcriptional networks may also explain why some individuals with autism or schizophrenia have considerable talent in certain cognitive domains, and the positive genetic correlation at the level of common variants between schizophrenia, autism and educational attainment.

We used genes identified for educational attainment as a proxy for intelligence and cognition. While there is not a perfect correlation between educational attainment and IQ, the educational attainment GWAS is the largest GWAS to date of a cognitive phenotype^24^. As such, it affords substantial power to robustly identify genes associated with cognition and intelligence. We reasoned that some cellular and molecular pathways will be preserved between educational attainment and general cognition. Indeed, we identified a modest enrichment for edu genes in the M_c_3 module that is also enriched for genes identified for cognition^29^. This module is conserved across different brain regions in humans and rodents. The M_c_3 module has been previously shown to be significantly enriched for genes associated with delayed recall and general fluid cognitive ability^29^. Both M_c_1 and M_c_3 are enriched for pathways involved in synaptic processes, in particular, the post-synaptic density and Reelin signalling pathway^29^. To underscore the specificity of this enrichment, we did not identify any enrichment for genes identified for height, a phenotype with large sample sizes and relatively well powered GWAS, but that has limited neural basis.

Our results also provide preliminary evidence that general cognition and educational attainment converge in common co-expression modules derived from normal brain and disease tissue supporting previous genetic correlation analyses that show a high to modest correlations between educational attainment and different aspects of cognition^39^. However, while educational attainment is often regarded as a proxy phenotype for cognition, it can also reflect contributions from other factors such as ‘grit’^42^ and socio-economic status^43^. In addition, while general cognitive aptitude may be enriched across modules conserved across different species of mammals, educational attainment can be thought of as a much more complex phenotype involving several uniquely human traits. Further, while some aspects of cognitive aptitude may primarily involve the hippocampus (such as working memory), educational attainment may involve many cortical areas. As such, it is possible that edu genes may be enriched in additional cortical networks that are not necessarily conserved across species.

It is difficult to tease apart the directionality in gene regulation from genome-wide association studies. Indeed, it is likely that these genes exhibit complex regulatory patterns with changes in specific developmental time point in specific tissues contributing to higher cognitive aptitude. Understanding how the expression edu genes changes in relation to educational aptitude will help understand if their co-expression with schizophrenia and autism risk genes lead to cognitive deficits, or conversely to talent. In this study, we have used the largest available datasets to identify both edu genes and modules associated with autism and schizophrenia, though, the autism co-expression modules were constructed using only about a third of the samples used for schizophrenia. Replication in a larger dataset will help confirm the validity of these findings and may aid in identifying both common pathways and directionality between the three phenotypes.

## Acknowledgments

VW was funded by the Cambridge Trusts and St. John’s College, Cambridge. RB was funded by The Pinsent Darwin Trust and the MRC fellowship. This work was funded by the Autism Research Trust and the Templeton World Charity Foundation, Inc. The research was carried out in association with the National Institute for Health Research (NIHR) Collaboration for Leadership in Applied Health Research and Care East of England at Cambridgeshire and Peterborough NHS Foundation Trust. The views expressed are those of the author(s) and not necessarily those of the NHS, the NIHR or the Department of Health.

## Author contributions

All authors designed the study. VW and RAIB analysed the data. VW and RAIB wrote the manuscript that was then edited by DG and SBC.

